# An optimized contact map for GōMartini 3 enabling conformational changes in protein assemblies

**DOI:** 10.1101/2025.11.14.688437

**Authors:** Gustavo E. Olivos-Ramirez, Luis F. Cofas-Vargas, Siewert J. Marrink, Adolfo B. Poma

## Abstract

Advances in structural biology, particularly cryo-electron microscopy, have enabled high-resolution characterization of complex biomolecular assemblies. These developments emphasize the need for computational approaches capable of describing biologically relevant conformational changes over extended timescales. GōMartini 3 is a coarse-grained approach that demonstrates computational efficiency and versatility across several systems, from protein-binding membranes and soluble proteins to intrinsically disordered proteins, while preserving key physicochemical features. In this work, we introduce an optimized approach that integrates dynamic contact information from AA-MD simulations to refine the contact map in GōMartini simulations. Specifically, we define high-frequency contacts (HFC), which reduce the number of original Gō contacts set by ≈20–30%, thereby improving the representation of conformational states beyond the original approach. Benchmarking different contact-selection criteria revealed that including intra- and interchain HFC captures structural flexibility and domain dynamics. The method was tested on three small soluble proteins and on the SARS-CoV-2 spike protein. Overall, the optimized contact map improves sampling efficiency and expands the accessible conformational landscape relative to the original GōMartini 3 implementation. The full framework is available as an open-source resource for large-scale simulations of biomolecular assemblies.

## Introduction

With the advancement of experimental techniques such as cryo-electron microscopy, it is now possible to solve structures of increasingly complex biological systems, often in the megadalton (MDa) range. ^1^ Experimental structures provide static or ensemble snapshots, but molecular dynamics (MD) simulations are essential to reveal the continuous conformational transitions underlying biological function. All-atom (AA) MD simulations provide atomic-level detail; however, their computational cost limits accessible system sizes and timescales, typically reaching only from nanoseconds to a few microseconds. Performing MD simulations at the millisecond timescale^2^ or with very large systems^3–6^ requires extensive high-performance computing resources, which are not always accessible. To overcome these limitations, coarse-grained (CG) approaches have emerged as an alternative, offering significant reductions in computational cost while enabling the exploration of large systems and longer timescales. ^7^

CG models reduce molecular complexity by mapping groups of atoms onto interaction sites or beads, thereby smoothing the energy landscape while maintaining a molecular-level representation.^8^ This simplification accelerates simulations by two to three orders of magnitude compared to atomistic models,^9^ allowing the observation of rare events such as folding, binding, and large-scale conformational transitions. Among CG frameworks, the Martini force field^10^ is one of the most widely used. Initially developed for lipid membranes, it has since evolved into a general-purpose model applicable to proteins, nucleic acids, and large supramolecular complexes, enabling simulations that reach the millisecond regime. ^11–14^ The latest Martini 3 release includes several approaches aimed at stabilizing protein structures: the elastic network (EN) to maintain structural integrity,^15^ the GōMartini 3 model that integrates native contact information for balanced flexibility,^16,17^ and the OLIVES multiscale pipeline which uses hydrogen-bonding information to stabilize the structure.^18^ Moreover, a recent approach termed as OliGōmers^19^ has been designed for multimer systems to study oligomerization and aggregation phenomena.

The GōMartini 3 framework is widely used to study single proteins and protein complexes, providing a balance between structural stability and conformational flexibility.^20–22^ This approach integrates Martini 3 nonbonded interactions with a Gō-like^23,24^ potential defined over native contacts (NCs) that are derived from a reference structure. The native contact map (CM) is typically constructed using geometric overlap (OV) and repulsive Contacts of Structural Units (rCSU) criteria,^25^ and Lennard–Jones (LJ) potentials are applied to stabilize the corresponding tertiary and quaternary interactions during simulations. However, because the CM is derived from a single static model (i.e., crystal structure), the resulting potential confines the system within one free-energy basin, restricting transitions between alternative conformational states. This limitation becomes especially critical for large or multichain complexes, where fixed native contacts may artificially constrain inter-domain motions or slow down conformational changes. ^17^ The switching^26^ and multiple-basin^27^ GōMartini variants partially address this issue by interpolating between CM derived from two known states, yet they require the availability of experimentally solved conformations for both. An alternative perturbation-based optimization approach (*PoGō*) showed to fine-tuned the *ɛ* and *σ* parameters in the Gō potentials using AA-MD data, capturing the essential dynamics for protein monomers as in AA-MD.^17,28^ For systems where only one structure is available, it remains challenging to model long-timescale conformational changes.

Multiscale modeling integrates two or more simulation approaches to bridge different spatial and temporal resolutions. ^29^ In biomolecular systems, such methods connect atomistic accuracy with CG efficiency, enabling the study of processes that span from nanometers to hundreds of nanometers and from nanoseconds to milliseconds. Multiscale approaches can generally be divided into two categories: hybrid (parallel) and sequential (serial). ^30,31^ In hybrid schemes, multiple levels of resolution coexist within a single simulation, for example, QM/MM or MM/CG, while sequential approaches use information from higher-resolution models to parameterize or guide lower-resolution representations. By combining the structural detail of atomistic simulations with the efficiency of CG models, multiscale strategies provide improved physical realism, access to extended timescales, and a substantial reduction in computational cost. ^32^ In this context, AA-MD simulations on the nanosecond-to-microsecond scale^33^ can be used to refine GōMartini 3 contact definitions, yielding a data-driven model that retains structural stability while enhancing conformational sampling. However, to achieve this integration, a dedicated framework is required to accurately translate atomistic dynamic information into the CG representation. This strategy benefits from and aligns with the FAIR principles,^34^ as it promotes the reuse of existing data to improve the accuracy of lower-resolution models.

In this study, we present a framework to optimize the CM for GōMartini 3 with the main objective of enhancing conformational sampling in large protein assemblies. Our approach (Figure 1) reduces the number of Gō contacts by constructing a refined contact map that includes only HFC derived from AA-MD simulations. We validated the HFC concept using three small soluble proteins (PDB IDs: 1UBQ, 1AOH, and 3W0K) and the SARS-CoV-2 S protein. Overall, the enhanced GōMartini 3 framework substantially improves conformational sampling by reducing native contacts, producing a more flexible and realistic CG model capable of exploring functionally relevant states across extended timescales.

**Figure 1:**
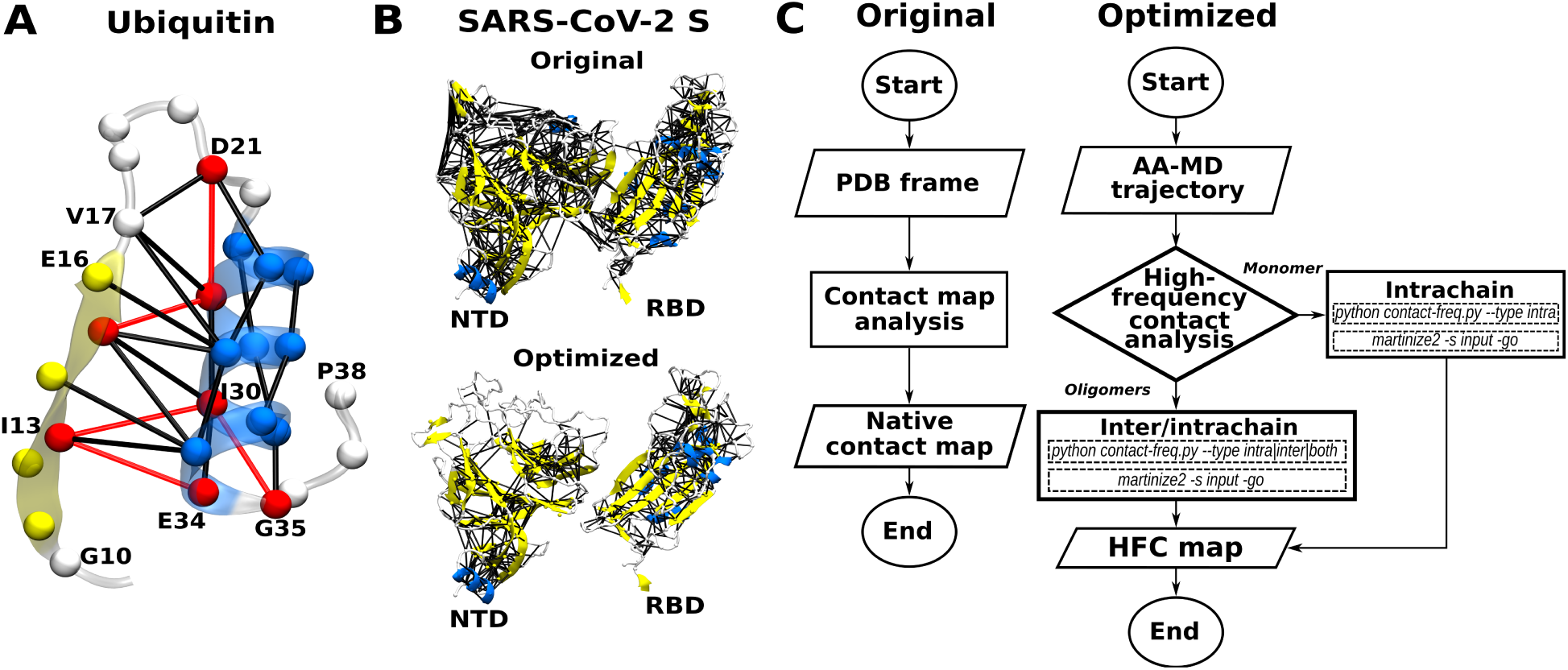
Refinement of HFC for GōMartini 3 simulations. (A) Residues 11–37 of ubiquitin, showing HFC (black lines) and low-frequency contacts (red lines) derived from atomistic MD simulations (1 *µ*s total sampling across five replicas). HFC are defined as residue pairs that persist in at least 70% of simulation frames. (B) Contacts between the receptor-binding domain (RBD) and the N-terminal domain (NTD) of the SARS-CoV-2 spike protein (PDB ID: 6VSB). The upper panel shows native contacts derived from the crystal structure, while the lower panel displays HFC extracted from 320 ns of atomistic MD simulations (five replicas). C) Comparison between the original algorithm for obtaining the contact map (left) and the optimization using HFC (right).

## Results and discussion

In Figure 1, we introduce the concept of high frequency contacts (HFC) used for enhancing GōMartini 3 CM. In the original workflow (Figure 1C), contacts are mapped from the crystal structure using the OV + rCSU concepts over residue pairs (i, |*i*− *j*| ≥ 3), resulting in a CM that preserves the folded topology. However, in the cases of protein assemblies, the high surface area in contact between protomers can generate excessive definitions of Gō contacts. This effect can overconstrain the system to a single energetic basin, limiting the conformational sampling. To address this limitation, our approach selectively corrects both inter- and intrachain contacts to reduce redundancy and enhance sampling. Specifically, we refine the CM by removing low-frequency contacts identified from AA-MD. As shown in Figure 1A, high-frequency (black) and low-frequency (red) contacts are overlaid for ubiquitin. The removal of low-frequency contacts, such as the hydrophobic pair I13–I30 (red lines in Figure 1A), still preserves the integrity of the hydrophobic core. This refinement becomes even more critical in complex systems such as the SARS-CoV-2 spike (S) protein. As shown in Figure 1B, the contact network illustrates the differences between the original and enhanced CM. By pruning low-frequency contacts, the coarse-grained (CG) model gains the flexibility required to explore multiple conformational states.

For validation, we benchmarked HFC in three small proteins (ubiquitin, cohesin domain, and glycoside hydrolase) and evaluated their stability against AA-MD simulations. Afterwards, using the SARS-CoV-2 S protein as a case study, we generated three distinct HFC maps for GōMartini 3 simulations (termed as optimized-1, -2, and -3) with the objective of determining the intra- and interchain contact effect. Our method quantifies both intra-and interchain contacts throughout the trajectory, defining an HFC as any contact with a frequency of ≥ 0.7. In addition, it identifies and selects the frame containing the highest number of HFC. In optimized-1, both inter- and intrachain HFCs defined in the selected frame, together with additional HFC identified throughout the AA-MD trajectory, were included (NCs = 5408). In optimized-2, the interchain HFCs identified in the selected frame were incorporated, while preserving the existing intrachain contacts from that same frame (NCs = 6973). Finally, in optimized-3, both inter- and intrachain HFC (NCs = 4966) were incorporated (see Methods and Table S1). We demonstrate that both intra- and interchain HFC enable the observation of significant conformational transitions, in contrast to the traditional approach. This implementation enables the study of large macromolecular assemblies with a more accurate and dynamic definition of native contacts, preserving tertiary and quaternary structure while facilitating efficient exploration of long-timescale conformational dynamics.

### Contact map optimization in small systems

We first validated the HFC refinement protocol using small protein systems. AA-MD simulations of 1 *µ*s each (five replicas per system) were performed for ubiquitin (PDB ID: 1UBQ),^35^ the cohesin domain (PDB ID: 1AOH),^36^ and glycoside hydrolase (PDB ID: 3W0K). HFC were extracted from these trajectories to construct refined CM, which were subsequently employed in GōMartini 3 simulations of equivalent length and replica count. Conventional GōMartini 3 simulations using crystal-derived CM were performed in parallel as controls. Compared to the crystal-derived CM, the refined HFC maps retained 80.4%, 72.1%, and 75.0% of contacts for 1UBQ, 1AOH, and 3W0K, respectively. This corresponds to an approximate 20–28% reduction in contact count relative to the standard protocol, yielding a less constrained model expected to better capture intrinsic protein flexibility.

To evaluate the effect of removing low-frequency contacts on model accuracy, we compared the RMSD and RMSF profiles obtained from AA-MD, the standard GōMartini 3 model based on the crystal-derived CM, and the enhanced GōMartini 3 model incorporating HFC. In addition, the Hellinger distance (H; ranging from 0 to 1)^37^ was used to quantify the similarity between RMSF distributions, where lower values indicate greater similarity. Empirically, values below 0.3 denote excellent agreement, while those up to approximately 0.7 still reflect acceptable similarity. For ubiquitin, the RMSD profile of the HFC model overlapped the AA-MD reference, while the RMSF profiles remained consistent (Figure 2A-C). Only residues 30–40 and 44–46 exhibited slightly increased flexibility (≈1.3 Å). RMSF distributions largely overlapped the atomistic reference, with an H = 0.34 for the crystal and H = 0.43 for the HFC. For the cohesin domain, the HFC model showed a small RMSD increase (≈1.0 Å) but achieved excellent agreement in flexibility, with H = 0.29 for the crystal and H = 0.31 for the HFC model (Figure 2D-F). In the glycoside hydrolase, the HFC produced a moderate RMSD increase (≈1.5 Å) while maintaining similar RMSF patterns, with distributional overlap slightly favoring the crystal (H = 0.20) over HFC (H = 0.43) (Figure 2G-I). Overall, the removal of low-frequency contacts still preserved structural stability and flexibility across all three systems. Nevertheless, for small globular proteins, the standard GōMartini 3 model remains slightly more consistent with AA-MD. Individual profiles for those systems are found in Figure S1-S3.

**Figure 2:**
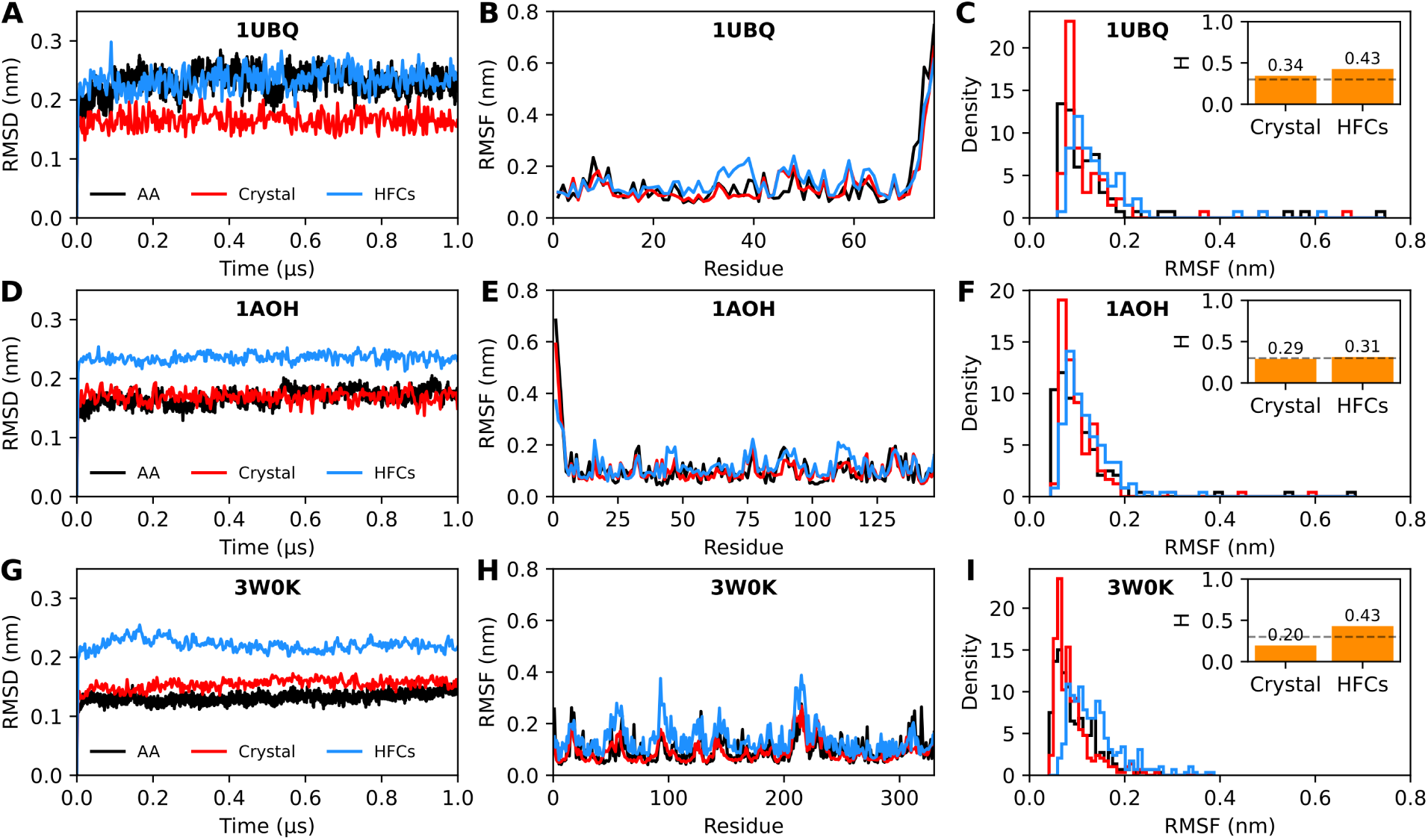
Flexibility of small proteins assessed by AA and CG simulations. Structural stability and flexibility of ubiquitin (1UBQ), the cohesin domain (1AOH), and glycoside hydrolase (3W0K) were evaluated from 1 *µ*s of atomistic MD (five replicas) and corresponding GōMartini 3 simulations using *ε* = 15.0 kJ/mol. Panels A–C show the results for 1UBQ, including RMSD, RMSF, RMSF distribution, and the estimation of the Hellinger distance (H) between the classical GōMartini 3 model (crystal-based contacts) and the enhanced model using HFC, both compared to the atomistic reference. Panels D–F and G–I show the same analyses for 1AOH and 3W0K, respectively. Dashed lines mark the critical H threshold used to quantify agreement between CG and AA distributions.

### Contact map optimization in protein assemblies

For oligomeric cases, such as the ≈0.6 MDa trimeric SARS-CoV-2 S protein, we evaluated the contribution of inter- and intrachain HFC to the stability and dynamics. Three HFC optimizations (optimized-1, -2, and -3) were constructed and validated against AA-MD simulations. These comparative setups (Table S1) were designed to identify the optimization scheme that best preserves structural stability while providing sufficient flexibility to enhance conformational sampling.

A detailed analysis of the principal domains (RBD, NTD, S1/S2, FP, and HR1) in the chain with the RBD-up revealed similar flexibility patterns for optimized-1 and optimized-3 (Figure 3A). The RMSF profiles of these two models showed enhanced mobility in the NTD, with optimized-3 most closely resembling the reference AA model. The RMSF of the RBD remained generally consistent across all models, while optimized-3 better reproduced the reference behavior in the S1/S2 and HR1 regions. RMSF of FP was also more accurately captured by optimized-3. For optimized-2, all domains displayed lower RMSF values, reflecting a more rigid behavior. Overall, these RMSF profiles indicate that optimized-1 and optimized-3 preserve the dynamic features of the AA model. The RMSD analysis further supported these observations (Figure 3B). All three optimized models reached RMSD convergence; however, optimized-2 exhibited lower values, indicative of reduced conformational exploration caused by its stiffer intrachain network. Optimized-1 and optimized-3, in turn, presented similar RMSD profiles, slightly higher than the reference model (gray dashed line in Figure 3B) but differing by only ≈1 . Moreover, H values between the RMSF distributions remained below 0.3 across all models (Figure 3C), confirming good agreement with the reference. Minor variations in H originated from local differences in the main domains (i.e., RBD and NTD) highlighted in the RMSF profiles as shadows. These results demonstrate that incorporating inter- and intrachain HFC preserves the flexibility of the GōMartini 3 models relative to the AA-MD simulation. Individual profiles for the S protein can be found in Figure S4-S5.

**Figure 3:**
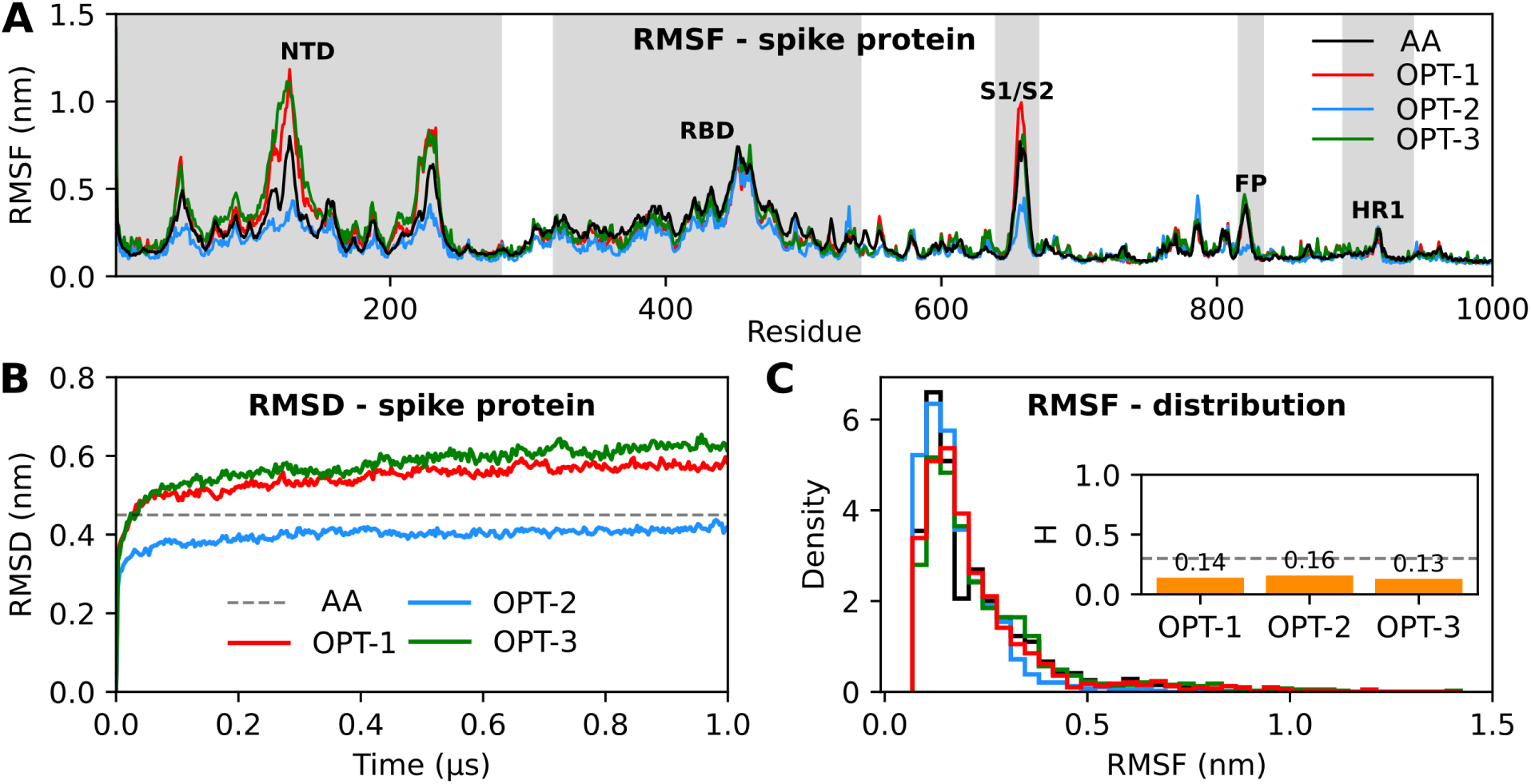
Flexibility of the SARS-CoV-2 spike protein in optimized GōMartini 3 simulations. Structural stability and flexibility of the trimeric spike protein were evaluated from 1 *µ*s simulations (five replicas) using three optimized GōMartini 3 configurations with *ε* = 15.0 kJ/mol. (A) RMSF profiles of the chain in the RBD-up conformation for the atomistic and optimized models. Shaded regions highlight the principal functional domains of the spike protein. (B) RMSD profiles for each model. The black line represents the average RMSD value (0.45 nm) from the atomistic reference model obtained from 320 ns of simulations (five replicas). (C) RMSF distributions and Hellinger distance (H) values for each optimized model compared to the atomistic reference. Dashed lines indicate the critical H threshold used to assess distributional similarity.

### Application in large-scale protein system

To further evaluate the robustness of the enhanced GōMartini 3 model, we extended the S protein simulations to a timescale of 10 *µ*s (10 replicas). Since the initial structure contained one RBD-up, we selected standard CVs to characterize its conformational fluctuations within the sampled space. Specifically, we measured the distance (nm) between the centers of mass of residues I338–W442 and Q512–G532 from the RBD-up and its opposite RBD, corresponding to the direction of the up-to-down transition, as well as the angular fluctuation (*θ*) of the RBD relative to a vertical vector defined by the HR1 domain. These CVs, previously employed to characterize RBD transitions,^38^ were used here to construct the free-energy landscape (FEL) and to compute the probability distribution of metastable states, as shown in Figure 4. Conformational states were identified by the ML-DBSCAN method^39^ (Figure S6).

**Figure 4:**
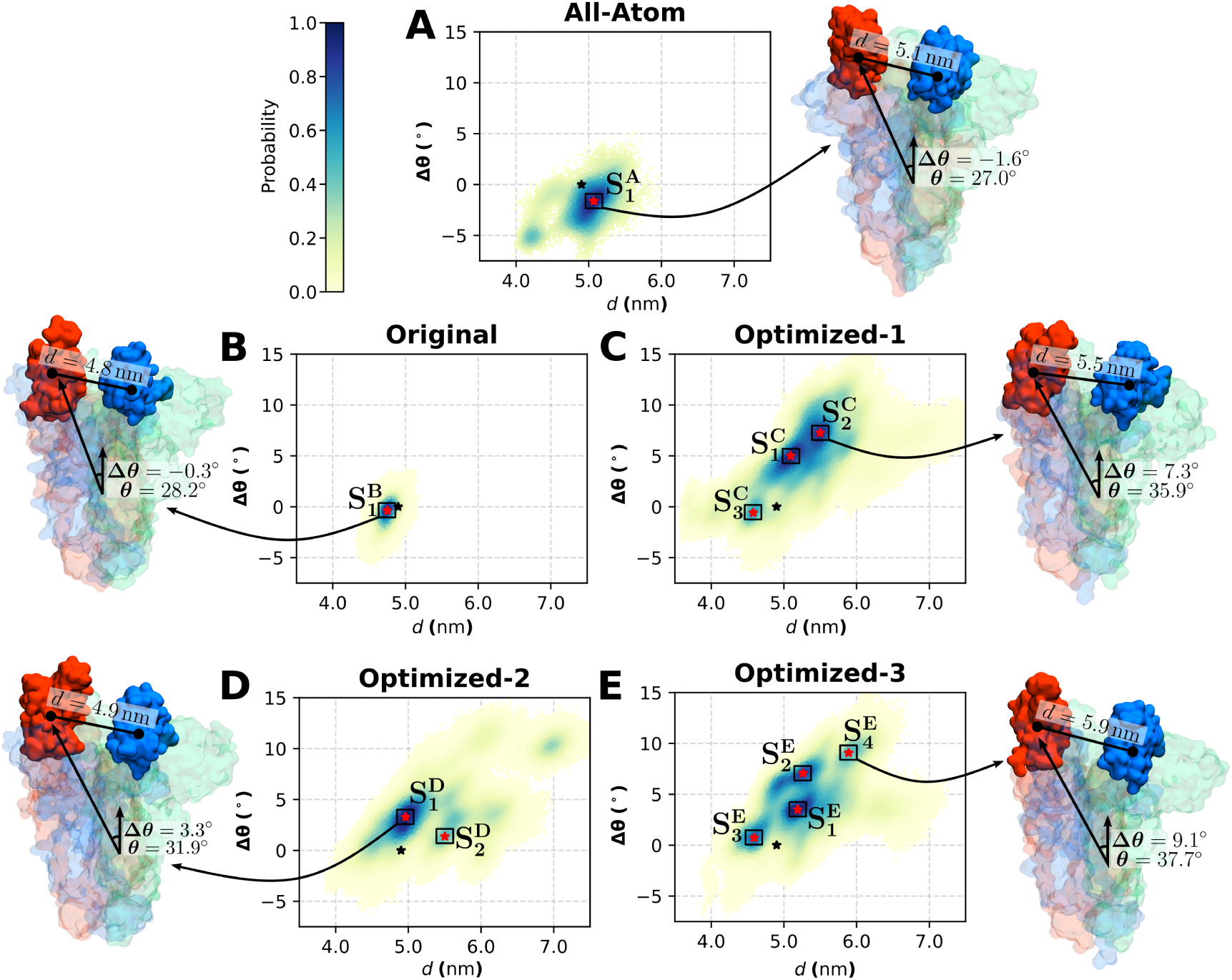
Probability states of the WT SARS-CoV-2 spike protein determined from 10 *µ*s of GōMartini 3 simulations (ten replicas) using *ε* = 15.0 kJ/mol. Different CM optimizations (1, 2, and 3) are compared with (A) the AA reference model and (B) the standard GōMartini 3. Panels C–E correspond to the three optimized models incorporating HFC (frequency > 70%). Red stars denote independent clusters (S*_n_*) and black ones the reference AA structure. The highest probability of each cluster is indicated for each metastate. Only clusters with probability ≥50% are presented. Snapshots indicate the more diverse centroid w.r.t. AA model. Distance (*d*) and Δ*θ* according to FEL are reported (Δ*θ* = *θ* − *θ*_0_; *θ*_0_ = 28.6*^◦^*).

The AA model was characterized by two well-defined minimas within the conformational space of the RBD-up. However, *S*_1_ concentrate most of the probability, with the second state presenting low probability (P*<*0.5) (Figure 4A). Although the AA-MD simulation was relatively short (320 ns), the RBD domain exhibited some flexibility, within the range of 4.0-5.5 nm of distance variation and angle fluctuation in the range of -7*^◦^* and -4*^◦^*. In contrast, the standard GōMartini 3 simulation (original) sampled a narrower conformational space, exhibiting a single dominant state (Figure 4B). This FEL was notably more confined, despite the extended simulation time and multiple replicas employed. Here, the RBD remained close to its original conformation (black start). The limited sampling capacity of the original model originates from an excessive number of Gō contacts within the RBDs, NTDs, and HR1 domains. This effect is especially relevant in oligomeric systems, where interfacial complexity increases the number of contacts and, consequently, the structural constraints that hinder conformational transitions.

In the three optimized models, the sampling space was expanded (Figure 4C–E). In these cases, the distance between the RBDs changed approximately from 3.6 nm to 7.5 nm, and the angle fluctuated between -4.9°and 15°(Figures 4C–E). Although all optimized models enhanced the sampling capacity, their main differences arose from the distinct probabilities assigned to each metastable state (Table 1). Optimized-1 presented three metastable states, optimized-2 presented two states, and optimized-3 presented four states, all of them with P > 0.5. Notably, optimized-2 concentrated the density in S_1_ and S_2_, while other minima were observed but with lower probabilities (P < 0.5). This suggests that optimizing only the interchain contacts is insufficient to improve the model, as intrachain interactions also play a key role in maintaining flexibility and enabling conformational transitions (Figure 4D). Meanwhile, optimized-1 and -3 presented higher probabilities distributed among three and four states, respectively, with similar coverage in the CV space. However, optimized-3 outperformed the other models by exhibiting multiple high-probability states (P > 0.5) spanning different distances and angles (Figure 4E).This optimization relied on fewer stabilizing contacts yet still improved conformational sampling while preserving structural stability.

**Table 1:** Number of interchain contacts in the metastable states for the optimized models.

| Model | CM | States | $P$ | Total NCs | ΔNCs | RBD NCs | Identity (%) |
| --- | --- | --- | --- | --- | --- | --- | --- |
| All-atom | — | $S_0$ | | $7144 \pm 46$ | — | $40 \pm 5$ | — |
| Original | 7289 | $S_0$ | | $6772 \pm 45$ | $\downarrow 372$ | $49 \pm 7$ | 33.8 |
| Optimized-1 | 5408 | $S_0$ | $<0.01$ | $6893 \pm 47$ | $\downarrow 251$ | $50 \pm 12$ | 21.5 |
| Optimized-2 | 6973 | $S_0$ | | $6838 \pm 72$ | $\downarrow 251$ | $55 \pm 10$ | 29.2 |
| Optimized-3 | 4966 | $S_0$ | | $6888 \pm 81$ | $\downarrow 256$ | $59 \pm 14$ | 30.8 |
| All-atom | — | $S_1^A$ | 1.0 | $7131 \pm 50$ | — | $39 \pm 8$ | — |
| Original | 7289 | $S_1^B$ | 1.0 | $6774 \pm 49$ | $\downarrow 515$ | $52 \pm 7$ | 80.5 |
| Optimized-1 | 5408 | $S_1^C$ | 1.0 | $6891 \pm 51$ | $\uparrow 1483$ | $45 \pm 12$ | — |
| | | $S_2^C$ | 0.93 | $6757 \pm 58$ | $\uparrow 1349$ | $55 \pm 8$ | — |
| | | $S_3^C$ | 0.61 | $6860 \pm 62$ | $\uparrow 1452$ | $38 \pm 10$ | — |
| Optimized-2 | 6973 | $S_1^D$ | 1.0 | $6757 \pm 58$ | $\downarrow 216$ | $32 \pm 15$ | — |
| | | $S_2^D$ | 0.50 | $6802 \pm 57$ | $\downarrow 171$ | $19 \pm 5$ | — |
| Optimized-3 | 4966 | $S_1^E$ | 1.0 | $6917 \pm 78$ | $\uparrow 1951$ | $52 \pm 13$ | — |
| | | $S_2^E$ | 0.88 | $6904 \pm 50$ | $\uparrow 1938$ | $42 \pm 11$ | — |
| | | $S_3^E$ | 0.78 | $6861 \pm 48$ | $\uparrow 1895$ | $59 \pm 10$ | — |
| | | $S_4^E$ | 0.50 | $6885 \pm 56$ | $\uparrow 1919$ | $62 \pm 17$ | — |

CM = number of Gō contacts in each contact map. Different superscript letters in $S_n$ (A–E) indicate the corresponding model: A = All-atom, B = Original, C = Optimized-1, D = Optimized-2, and E = Optimized-3. $P$ = probability. $\Delta$ NCs = difference in contacts with respect to AA for $S_0$ and to the CM for the different states ( $S_n$ ). Identity = number of contacts equal to the AA for $S_0$ and to the CM for the different states ( $S_n$ ).

To further evaluate the model, we calculated the number of interchain contacts in each state. This analysis was performed for the highest-probability states and for the conformations closest to the initial state. The initial state is defined as S_0_ (black stars in Figure 4). Although S_0_ was not a high-probability state in the simulations with HFC, it was included because it provides insights into contact reorganization when the system explores those co-ordinates. For this purpose, all conformations (frames) within the CV range of the initial structure (RBD distance = 4.92 nm and angle = 0.0*^◦^*, with a standard deviation of ±0.01) and of the high-probability metastable states were included in the analysis. To determine Gō contacts, the selected frames were backmapped using the one-bead-per-residue approach. ^40^ The total number of interchain contacts in the spike trimer, as well as those specifically associated with the RBD-up, are summarized in Table 1.

The total number of contacts differed substantially among all GōMartini 3 simulations (ranging from 6772 to 6893) compared to the AA-MD simulation (7441 ± 46) in the S_0_ state. These discrepancies arise from the Gō potential and the Martini 3 force field, both of which aim to preserve structural stability. As a result, although the systems explore similar regions of the CV space, the overall assembly diverges in specific structural regions, leading to variations in the contact network. This effect is particularly evident in the RBDs, where the total number of contacts remains comparable across models, yet their identities differ markedly (Table 1). The overlap of RBD contact identities ranged from 21.5% to 33.8%. Theoretically, a highly accurate model should reproduce not only similar conformations but also similar contact identities. However, as previously discussed, the distinct configurations of each optimized model likely constrain conformational changes at specific sites, preventing identical contact rearrangements. An additional explanation may be that other CVs, beyond those used here, could better capture the transitions and improve consistency in the contact identity among optimized models.

Besides, the metastable states observed in the optimized models displayed distinct contact profiles. In optimized-1 and -3, the total number of contacts increased markedly, even though their CM were constructed exclusively from HFC (Table 1). In contrast, optimized-2 exhibited metastable states (*S^D^* and *S^D^*) with contact counts similar to those of the initial CM, indicating limited structural reorganization. Although the number of RBD contacts remained within the standard deviation across models, the reduced number of states and their associated probabilities suggest that optimized-2 was the least effective configuration. Optimized-1 showed improved conformational sampling with three distinct metastable states, yet optimized-3 further enhanced sampling and stability, displaying up to four high-probability states. In optimized-3, the total number of contacts increased substantially relative to the original CM. The minima *S^E^*, *S^E^*, *S^E^*, and *S^E^* exhibited an average increase of ≈1900 total contacts and elevated RBD contact counts of 52 ± 13, 42 ± 11, 59 ± 10, and 62 ± 10, respectively. These states occupied regions of the CV space with larger angle (*>* 5*^◦^*) and distance (*>* 5 nm) values compared to the initial configuration (Figure 4E). Notably,*^E^* corresponds to a more extended RBD-up conformation, consistent with experimentally observed structures (PDB IDs: 7SBL and 7UPY), underscoring the accuracy of this optimization. Overall, these findings demonstrate that optimized-3, despite employing the smallest amount of contact information (only 4966 HFC), consistently produced the largest number of high-probability minima. This indicates that the model effectively recovers key stabilizing contacts while achieving superior conformational sampling efficiency.

Further work is required to optimize the atomistic simulation time necessary to determine HFC. Since this study aims to develop an improved methodology for sampling systems in the MDa range, AA simulations for such large assemblies may still represent a major limitation. Our results indicate that incorporating only a few hundred nanoseconds of replicated AA trajectories can substantially enhance sampling efficiency. It is also worth noting that our work recycled trajectories from previously published simulations. In this way, the reuse of existing atomistic information is promoted as a practical strategy to improve CG models in specific cases, reducing both computational cost and data redundancy. Moreover, our current approach has focused exclusively on reducing the number of contacts defined in the CM. Recent work by Kalutskii et al. ^28^ has instead concentrated on refining the LJ potentials. In this sense, both concepts could be combined to improve not only the accuracy but also the sampling efficiency of CG simulations. Additionally, the implementation of a Gō-like network with potentials dependent on distance fluctuations extracted from AA simulations may represent another promising route for optimizing the GōMartini 3 model.

## Conclusion

This work implements an approach to optimize the contact map and enhance the GōMartini 3 framework. We introduce the concept of high-frequency contacts (HFC) for the simulation of large protein assemblies. Structural and stability analyses (i.e., RMSD and RMSF) performed on small globular proteins demonstrate that HFC preserves the native structure within the same range as the classical approach, which uses the experimental structure as a reference to build the GōMartini protein model. The classical method yields a closer description of the native state, with limited fluctuations outside it, closely resembling the behaviour of the reference all-atom model. In oligomeric systems such as the SARS-CoV-2 S protein, the inclusion of HFCs significantly improved conformational sampling around the native state. By benchmarking different optimized contact maps, we found that incorporating both inter- and intrachain HFC resulted in a broader exploration of the conformational landscape. This global optimization of native contacts was the most efficient strategy in terms of Gō contact information, enabling the sampling of a larger number of conformational states (up to four) with a minimal set of contacts. Moreover, these states were more evenly distributed across the conformational space.

Our implementation is particularly valuable when a wide exploration of the conformational landscape is required, such as for processes occurring on the millisecond timescale. Additionally, reducing the number of Gō contacts while still retaining the native state of protein assemblies provides a more realistic biophysical description of their response to mechanical forces or temperature perturbations. Our work can boost other recent implemen-tation such a Switching Gō-Martini and Multiple-Basin Gō-Martini implementations^26,27^ to explore the vast protein conformational space between different metastable (active and inactive) states. Similarly, our work can be combined with *PoG*Ψ*o* (an optimization of GōMartini 3 Lennard-Jones parameters^28^ for capturing the protein essential dynamics.

The contact map optimization protocol we developed uses the martinize2 tool and delivers new parameters compatible with GōMartini 3. Ultimately, this work may help identify previously inaccessible conformational states that underlie the chemical or biological functionality of complex biomolecular assemblies in Martini 3.

## Methods

### All-atom molecular dynamics simulations

To assess the applicability of HFC within the GōMartini 3 approach, we first evaluated the accuracy of the model using three small protein systems: ubiquitin, the single cohesin domain, and the glycoside hydrolase (comprising 76, 147, and 330 amino acids, respectively). The atomic structures of these proteins were retrieved from the Protein Data Bank^41^ (PDB IDs: 1UBQ, 1AOH, and 3W0K, respectively). The proteins were repaired using PDBFixer.^42^ Missing atoms were added and protonation was performed at pH = 7.4. AA-MD simulations were conducted in GROMACS 2023.5. ^43^ Each system was placed in a cubic box with 10 Å of padding. TIP3P water molecules were used for solvatation. The system was neutralized by adding sodium (Na^+^) and chloride (Cl^−^) counterions at a concentration of 0.15 M. Energy minimization was performed using the steepest descent algorithm^44^ for 50000 steps. Equilibration was performed in two stages: NVT (100 ps) and NPT (200 ps). Temperature was equilibrated at 300 K using the V-rescale thermostat,^45^ while pressure was equilibrated at 1 bar using the Parrinello–Rahman barostat. ^46^ During minimization and equilibration, harmonic restraints of 1000 kJ mol^−1^ nm^−2^ were applied to the C-alpha atoms in their Cartesian coordinates. The LINCS algorithm^47^ was used to constrain all bonds, and long-range electrostatics were treated with the particle-mesh Ewald (PME) method.^48^ Production simulations were conducted for 1 *µ*s with five independent replicas in the NPT ensemble, using an integration timestep of 2 fs with the leap-frog integrator.^49^

In addition, we studied the case of the WT S protein from SARS-CoV-2. Atomistic trajectories of the WT S protein with a single RBD-up were obtained from a previous work.^50^ In that study, the WT structure (PDB ID: 6VSB) was retrieved from the Protein Data Bank. MD simulations were performed using the Amber ff14SB force field.^51^ The system was placed in a box with a padding of 12 Å and solvated using the TIP3P water model.^52^ Sodium and chloride ions were added to neutralize the system at a concentration of 0.150 M.

Energy minimization was conducted using the steepest descent algorithm with 2000 steps, followed by conjugate gradient equilibration with 3000 steps. Temperature equilibration was performed in two steps: NVT heating from 0 to 100 K over 50 ps, followed by NPT heating from 100 to 300 K over 100 ps. Harmonic restraints of 10 kcal mol^−1^ Å^−2^ were applied during the minimization and equilibration steps. These restraints were gradually reduced from 10 to 0.1 kcal mol^−1^ Å^−2^ through a short 6 ns simulation at 300 K. Pressure equilibration was performed using the Monte Carlo (MC) barostat^53^ at 1 atm. Temperature control was achieved with the Langevin thermostat^54^ using a collision frequency of 1 ps^−1^. Hydrogen mass repartitioning (HMR) was applied to the systems to achieve integration time steps of 4 fs.^55^ Production was performed for 320 ns with five replicas (1.6 *µ*s in total). Based on the RMSD (Figure S1), we determined that this system stabilizes after 80 ns. For this reason, for the analysis of HFC, we only used the convergence phase (last 240 ns of each replicate). Finally, we concatenate all the trajectories (6000 frames in total).

### Contact map determination protocol

To explore the conformational space of the protein systems, we performed CG simulations using the GōMartini 3 approach.^11,16,17^ This requires the definition of a CM, which is generated through a combination of the OV and rCSU methods. ^25^ The CM is computed from an atomistic structure, and the resulting contacts are incorporated into the GōMartini 3 simulations as Lennard-Jones (LJ) potentials. This approach is widely adopted and has become a standard in CG simulations.^17^ To improve the capabilities of this methodology, a careful selection of the optimal configuration of contacts was done. For this, we tested optimized CMs based on a selection of the HFC. Figure 1C shows the step by step protocol developed for enhancing the GōMartini 3 approach, and Table S1 shows the definition of each optimized CM.

The first step involved converting each frame (from the AA-MD simulations) to PDB format, as the script used for CM calculation only supports this file type. A contact was defined when the distance between C*α* atoms of those residues was within 3–11 Å. Frequencies were calculated for each pair of contacts across all frames. Only contacts with a frequency ≥ 0.7 were selected. HFC were identified for both intra- and interchain interactions. After determining HFC, a reference frame was selected for GōMartini 3 simulations. The selected frame was the one containing the highest number of HFC, obtained by comparison between the list of HFC and the contacts of each individual frame. In addition, if a contact is listed as of high frequency but not present in the reference structure (due to longer distances), the average distance of those pairs over the trajectory was calculated and then used to compute the *r_min_* values and define the virtual sites. Finally, using martinize2 tool,? new topology files (ITP files and CG structure) were generated keeping the list of HFC, while the original contacts from the reference frame (i.e., low frequency contacts) were removed.

All these steps were automatized using a custom pipeline built around two Python scripts designed to streamline the identification and classification of HFC and the generation of input files. These scripts allowed us to identify HFC efficiently and build topologies for subsequent GōMartini 3 simulations. The first script, termed *traj_to_pdb.py*, splits a trajectory into individual PDB frames at user-defined intervals, controlled by the --stride parameter. During this step, the chain index ranges of interest must be specified with the --range parameter (i.e., 1–1121, 1122–2242, 2243–3364; for the S protein). The second script, termed *contact_freq.py*, processes each frame to determine HFC and classifies them as interchain, intrachain, or both, depending on the --type argument. Using these criterion, we constructed the corresponding CM for the three optimized models. When using --type both, the optional --all-hf flag can be enabled to include HFC that are present along the trajectory into the selected reference frame. Additionally, the --go-eps parameter allows customization of the epsilon value used in the Gō potential (default: 9.414 kJ/mol). For multichain (oligomeric) systems, chain IDs are specified using the --merge option. A contact-frequency threshold is set via the --threshold parameter, commonly defined as 0.7, meaning the contact must be present in at least 70% of the frames. If DSSP^56^ is not available in the system path, the path to the mkdssp executable must be explicitly defined using the --dssp option. Once the reference frame containing the highest number of HFC is identified, the corresponding contact map is passed to *martinize2* to generate GōMartini 3 topology files.

For the small protein system, we evaluated only HFC of intrachain type, as these were single protomers. For the case of the S protein, we evaluated three different CM configurations. (1) a first optimized CM including all HFC (5408), both inter- and intrachain contacts of the reference structure, and additional ones that are not found in that structure but are mapped along the AA trajectory. (2) a second optimized CM including only inter-chain HFC, while the intrachain part was treated as non-high frequency (6973). Similarly, the reference frame containing HFC was selected. (3) a third optimized CM including only HFC of the inter- and intrachain space (4966), using the reference frame containing HFC as well. Since HFC were used in 1-3, these were referred to as optimized- 1, -2 and -3, respectively. All the scripts used in this protocol are available at https://github.com/GoMartini3-tools/ContactFreq/tree/main. The interaction energy of all GōMartini 3 simulations was set to 15.0 kJ mol^−1^.

### GōMartini 3 simulations

Once each optimized set of NCs was defined, GōMartini 3 simulations were performed with GROMACS 2023.5^43^ using the Martini 3 force field.^11^ The system topology was built using the martinize2 algorithm^57^ and secondary structure was assigned using DSSP v3.1.4. The initial structure was subjected to vacuum energy minimization for 5000 steps using the steepest descent algorithm. The system was then solvated in a cubic box of 10 Å of padding for the small proteins and in a 12 × 12 × 25 nm^3^ box for the S protein, using Martini water beads. The systems were neutralized by adding sodium and chloride ions to reach a concentration of 0.15 M. A second energy minimization step was performed for the solvated and neutralized system for 5000 steps. During equilibration, position restraints were applied to backbone beads (BB). Equilibration was carried out in two stages. The first stage was performed in the NVT ensemble at 300 K using the V-rescale thermostat,^45^ with a coupling time of 1.0 ps. This phase lasted 5 ns with integration steps of 20 fs. The second equilibration step was carried out in the NPT ensemble at 1 atm using the C-rescale thermostat,^58^ with isotropic pressure coupling = 10^−4^ bar^−1^ and a pressure coupling time constant of 12 ps. This phase lasted 10 ns and it also used 20 fs integration time. Production was conducted under NPT. Five replicas of 1 *µ*s each (totaling 5 *µ*s) was performed intially for all the systems. Then, for the S protein, we conducted a 10-*µ*s MD simulation with 10 replicas (cumulative 100 *µ*s) for each optimized protocol and for the original CM.

### Identification of metastable states via clustering

To determine metastable states in our CV space (RBD distance and angles) we employed a cluster methodology based on the multi-level density-based spatial clustering of applications with noise (ML-DBSCAN) algorithm developed by Liu et al. ^39^ . This method is based on the original DBSCAN. ML-DBSCAN is a density-based clustering method that can effectively detect metastable states from massive MD conformations. In this analysis we included all data-points from the 10 *µ*s with the 10 replicas. Using DBSCAN, a mestastable state is found when a large number of neighbors is found within the high density. The ML-DBSCAN performs the clustering in different density levels and then combines the results to obtain the hierarchical clusters within the FEL. The analysis was performed first, discarding points that had less than a 0.1 probability in the FEL. CVs data points were normalized (z-scored) and then clustered across successive density levels using decreasing neighborhood radii (*ɛ* = 0.045-0.030) and a fixed minimum number of neighbors (min_samples = 600). The resulting clusters from each level were integrated into a coarse-to-fine hierarchy, where persistent states were consolidated based on their parent–child overlap (persistence threshold ≥0.7) and a minimum population of 1500 conformations. We used an adapted GPU-accelerated implementation based on RAPIDS cuml v.25, ^59^ preserving the scheme of Liu et al. ^39^ while enabling efficient processing of million-frame datasets.

## Author Contributions

G.E.O.R. and L.F.C.V. contributed equally to this work. G.E.O.R. performed MD simulations, contributed to the algorithm development, analyzed and validated the data, and drafted the manuscript. L.F.C.V. developed the ContactFreq algorithm, performed MD simulations, and contributed to manuscript revision. S.J.M. participated in discussions and provided critical feedback. A.B.P. conceived the study, supervised the work, contributed to the algorithm discussion, acquired funding, and revised the manuscript. All authors have given approval to the final version of the manuscript.

## Acknowledgement

A.B.P. acknowledges earlier discussions on the topic with Rodrigo A. Moreira, financial support from the National Science Center, Poland, under grant 2022/45/B/NZ1/02519. This work thanks the Polish high-performance computing infrastructure PLGrid (HPC Center: ACK Cyfronet AGH) for providing computer facilities and support within computational grant no. PLG/2024/017332 and PLG/2025/018510.

## Supporting Information Available

All input files, simulation data, and processed results are openly available at Zenodo (DOI: 10.5281/zenodo.16023027) and are also included within the Supplementary Material.

## TOC Graphic

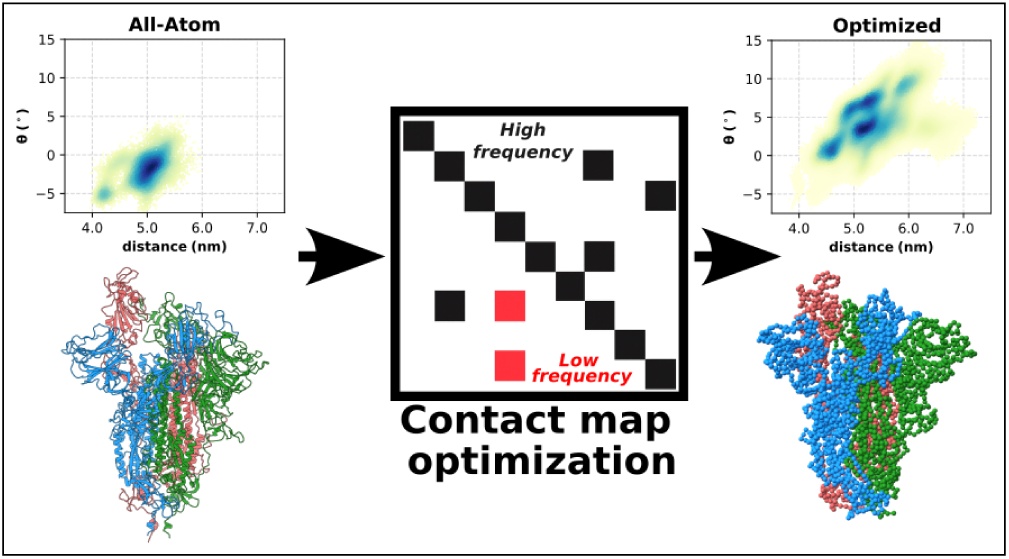

