## Supplementary Material for "An optimized contact map for GōMartini 3 enabling conformational changes in protein assemblies"

Table S1: Summary of the contact information used in each method.

| **Method** | **Contact information** |
|---|---|
| AA | None |
| Original | NC from the initial structure |
| Optimized-1 | HFC interchain + HFC intrachain + HFC that appear during the AA-MD |
| Optimized-2 | HFC interchain + intrachain contacts from the reference structure |
| Optimized-3 | HFC interchain + HFC intrachain |

NC = native contacts. HFC = high-frequency contacts.

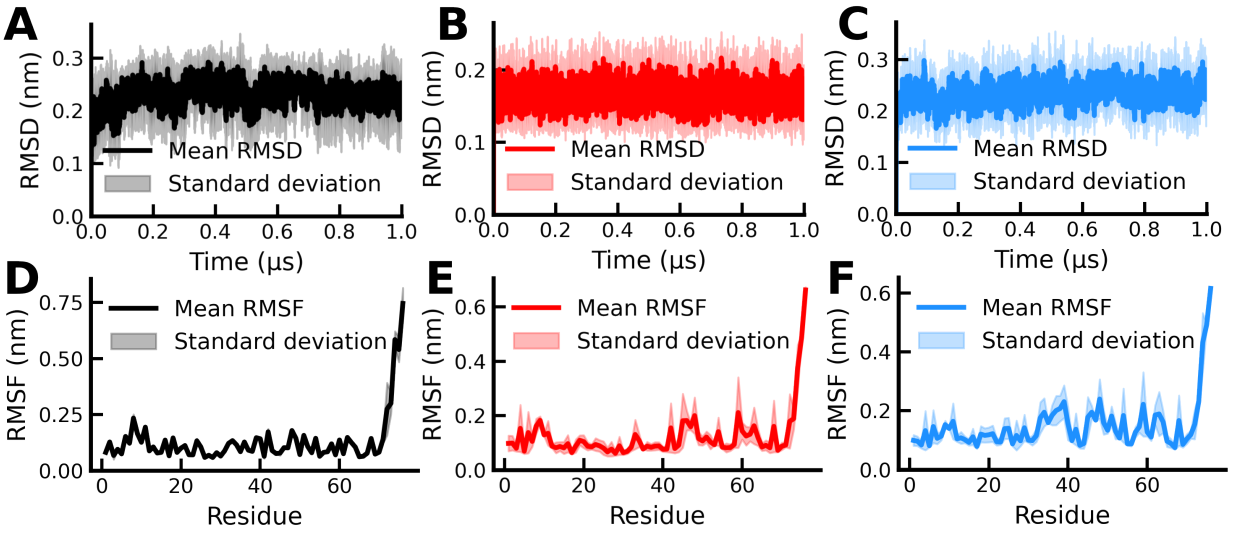


Figure S1: Time evolution of the average RMSD and RMSF for all simulation replicas (R1–R5) of the ubiquitin protein (PDB ID: 1UBQ), in AA-MD (black), GōMartini 3 with the crystal contact map (red), and GōMartini 3 with the HFC map (sky blue).

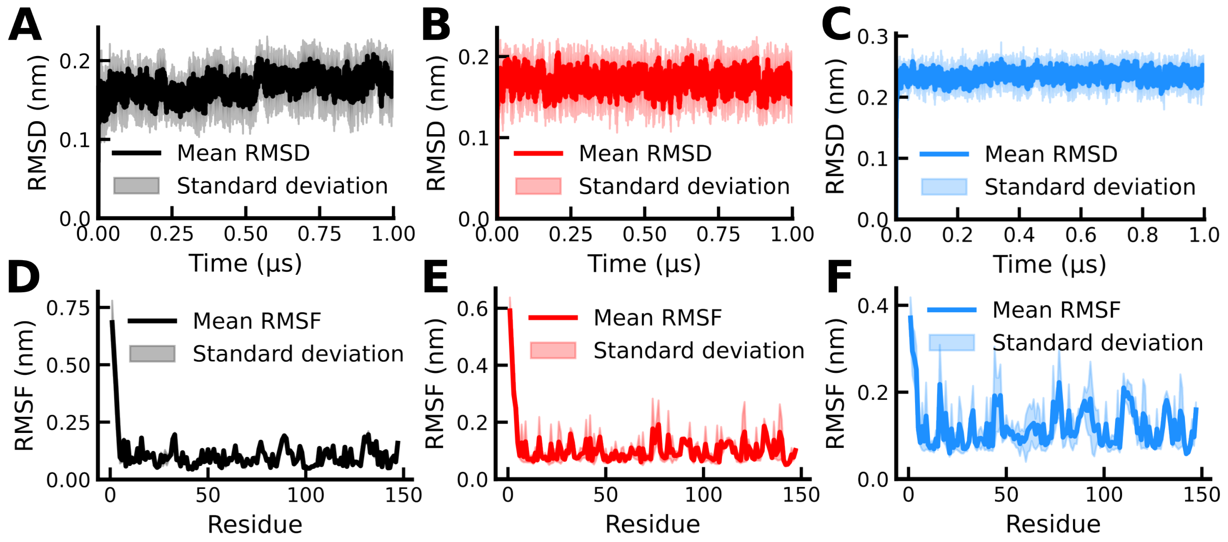


Figure S2: Time evolution of the average RMSD and RMSF for all simulation replicas (R1–R5) of the single cohesin domain protein (PDB ID: 1AOH), in AA-MD (black), GōMartini 3 with the crystal contact map (red), and GōMartini 3 with the HFC map (sky blue).

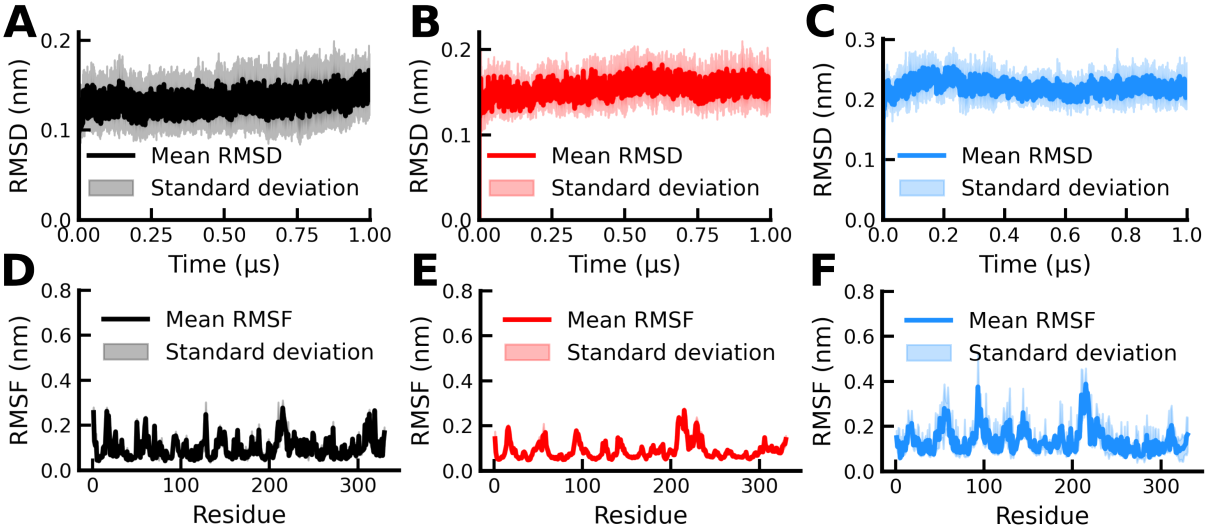


Figure S3: Time evolution of the average RMSD and RMSF for all simulation replicas (R1–R5) of the glycoside hydrolase protein (PDB ID: 3W0K), in AA-MD (black), GōMartini 3 with the crystal contact map (red), and GōMartini 3 with the HFC map (sky blue).

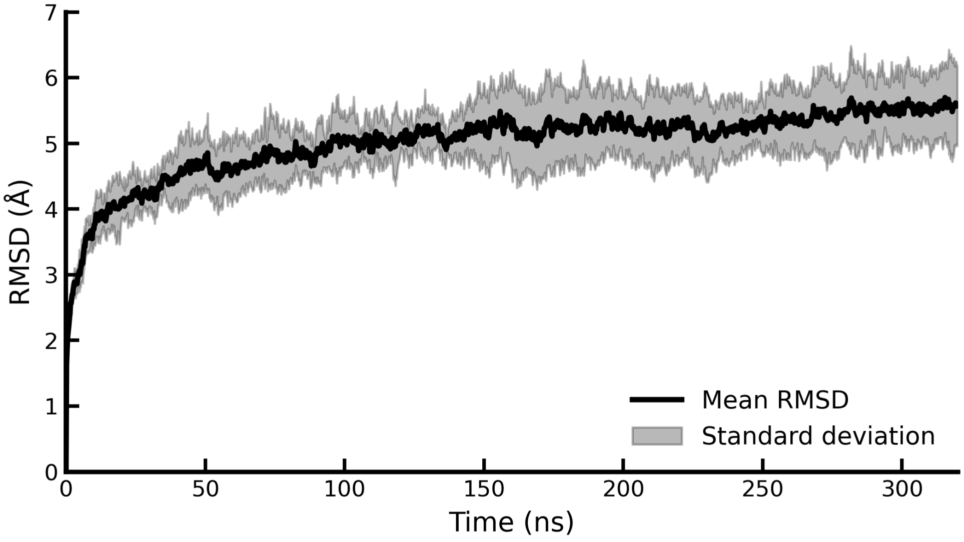


Figure S4: Time evolution of the average RMSD for all AA-MD simulation replicas (R1–R5) of the SARS-CoV-2 S protein.

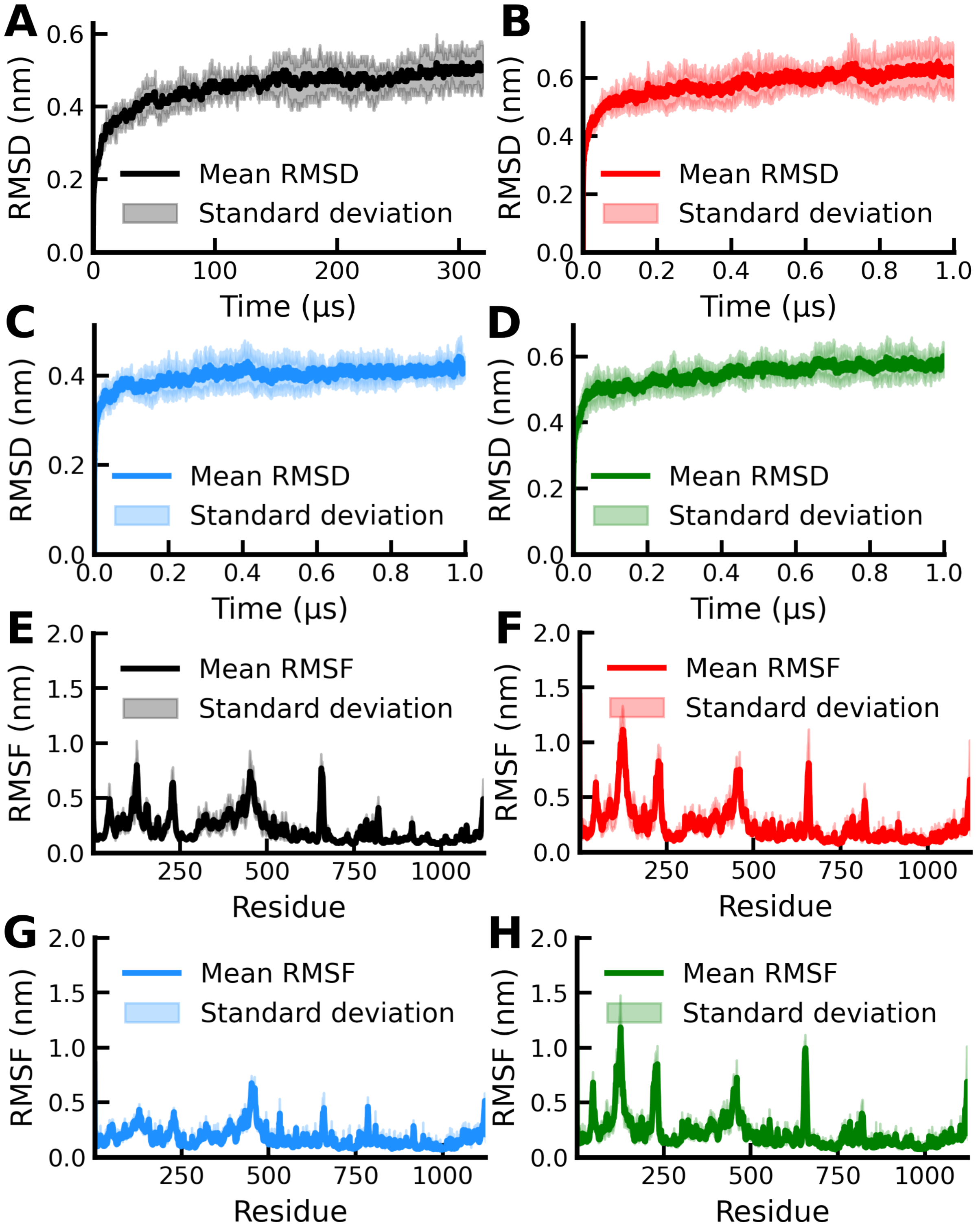


Figure S5: Time evolution of the average RMSD and RMSF for all simulation replicas (R1–R10) of the SARS-CoV-2 S protein (PDB ID: 6VSB), in AA-MD (black), GōMartini optimized-1 (red), optimized-2 (sky blue), and optimized-3 (green).

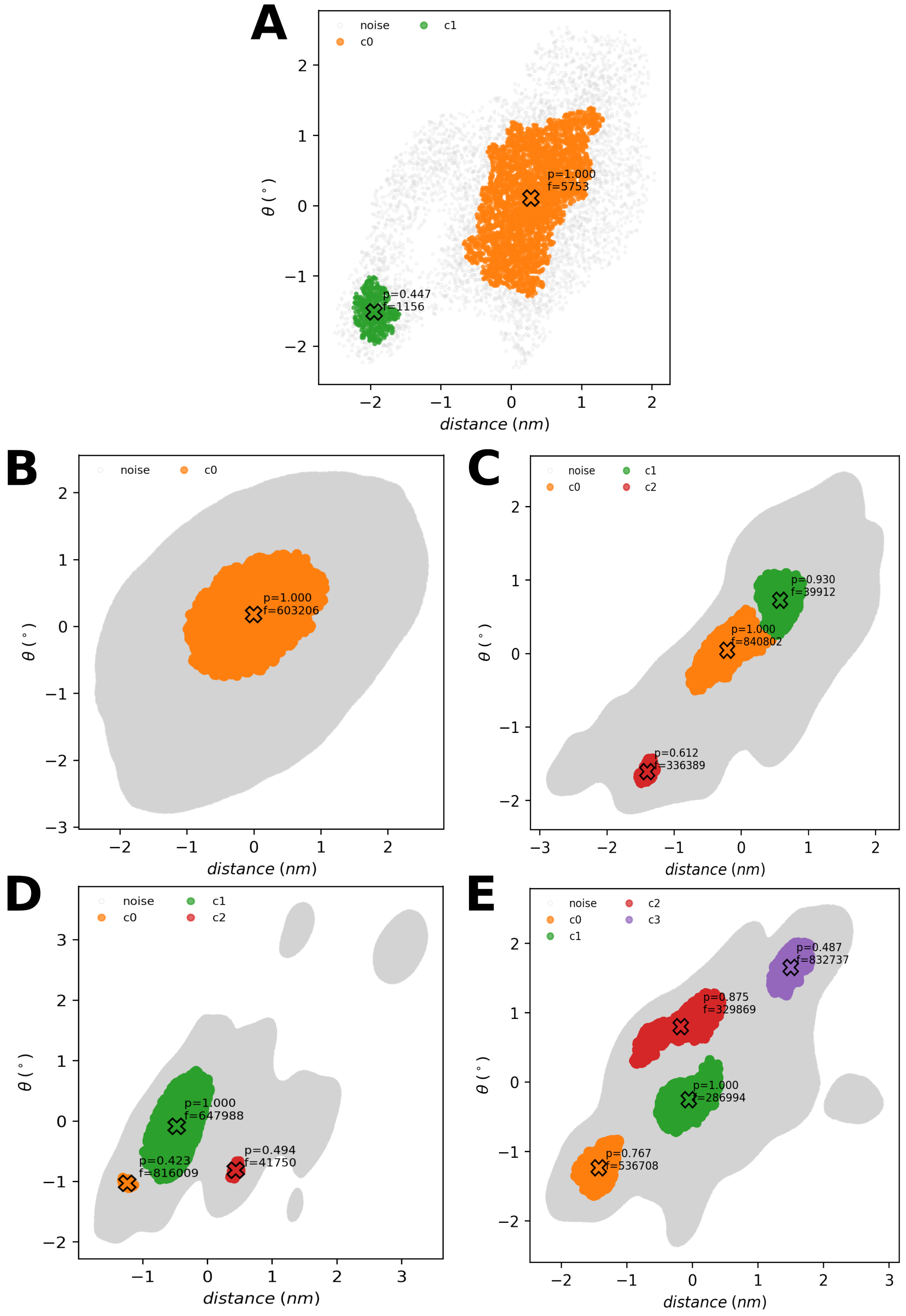


Figure S6: ML-DBSCAN of the most probable conformations of the SARS-CoV-2 S protein based on the CVs used in the FEL, from AA-MD (A), GōMartini 3 Original (B), Optimized-1 (C), Optimized-2 (D), and Optimized-3 (E) simulations.
